# Bacterial fucosidase enables the production of Bombay red blood cells

**DOI:** 10.1101/695213

**Authors:** Tiansheng Li, Juan Ye, Lei Wang, Lin Zou, Yameng Guo, Linlin Hou, Danfeng Shen, Xiaohong Cai, Haobo Huang, Guiqin Sun, Li Chen

**Affiliations:** Department of Medical Microbiology, Key Laboratory of Medical Molecular Virology of Ministries of Education and Health, School of Basic Medical Sciences, Fudan University, Shanghai 200032, China; Cancer Institute (Key Laboratory of Cancer Prevention and Intervention, China National Ministry of Education, Key Laboratory of Molecular Biology in Medical Sciences, Zhejiang Province, China), The Second Affiliated Hospital, School of Medicine, Zhejiang University, Hangzhou 310009, China; Key Laboratory of Cell Differentiation and Apoptosis of Ministry of Education, Department of Pathophysiology, Shanghai Jiao Tong University School of Medicine, Shanghai 200025, China; Blood Transfusion Department, Ruijin Hospital, Medical School of Shanghai Jiao Tong University, Shanghai 200025, China; Department of Blood Transfusion, Fujian Medical University Union Hospital, Gulou District, Fuzhou City, Fujian Province 350001, China; College of Medical Technology, Zhejiang Chinese Medical University, Hangzhou 310053, China

## Abstract

We present a method to produce H antigen-deficient red blood cells (RBCs) for transfusion to individuals with anti-H antibodies. A fucosidase from bacteria was heterologously expressed efficiently in E. coli and has been demonstrated to completely remove H antigens on the surface of human RBCs in a facile conversion process. The approach we describe here holds promise for making H-deficient RBCs available for a rare population beyond ABO types.

## Introduction

The ABO blood groups are vitally important in blood transfusion and organ transplantation^1,2^. Universal red blood cells (RBCs) are the ultimate goal to which scientists and researchers in the transfusion field have been dedicated for decades^3^. Previous studies demonstrated that A or B RBCs can be changed into enzyme-converted group O red blood cells (ECO RBCs) through treatment with α-N-acetylgalactosaminidase or α-galactosidase^4^, and the data of phase I and phase II clinical trials showed that B-ECO RBCs were safe and functional when transfused to O-type individuals^5,6^. Bacterial glycosidases have been identified to be capable of efficient removal of A and/or B antigens from human RBCs, with low consumption of recombinant enzymes^7,8^. This enzymatic technology provided a simple way to convert all donated blood into type O, which has great potential to eventually reduce blood shortages and make transfusions safer.

Beyond the ABO blood group, the H antigen, the precursor of A and B antigens, was found to be deficient on the surface of RBCs in a rare population, which is called the Bombay or Para-Bombay phenotype^9^. There are strong anti-H antibodies in the plasma of Bombay phenotype individuals, while weak anti-H antibodies are commonly observed in Para-Bombay individuals^10^. Transfusion to patients in the Bombay group requires full cross-compatibility, which means that only Bombay-type RBCs can be transfused to Bombay phenotype individuals^11^. However, transfusion to Para-Bombay individuals normally consists of A-, B- or AB-type RBCs, partially because of the lack of an H-deficient RBC supply for urgent blood transfusion^10^. Efforts have been made to remove H antigens from O-type blood to produce Bombay-type RBCs for transfusion to rare populations, but no efficient fucosidase has yet been identified^12,13^. Here, we report a bacterial fucosidase with exclusive substrate specificities for H blood group structures that can produce H-deficient RBCs, which can be used for transfusion to Bombay or Para-Bombay phenotype individuals with anti-H antibodies. This enzyme would make H antigen-deficient RBCs available in a cheap and simple way and create the possibility of safe transfusion for these rare communities.

## Results and Discussion

A panel of deduced fucosidase genes were identified in *Elizabethkingia meningoseptica*, including the first core α-1,3 fucosidase in our recent report^14,15^. One of these genes was found to encode a functional fucosidase with high enzymatic activity against α-1,2 fucosylated substrates. The recombinant enzyme was heterologously expressed efficiently without its N-terminal signal and putative transmembrane sequence (amino acid 35-775) with a yield of 1 g/L (designated csFase I). csFase I has a broad pH optimum between 5 and 7 (**Supplementary Figure 1 a)** and shows its highest activity at 37°C (**Supplementary Figure 1 b**). The substrate specificity result demonstrated its exclusive fucosidase activity without other detectable exoglycosidase features (**Supplementary Figure 1 c**). We also determined the linkage specificity of csFase I with a panel of oligosaccharides that contain representative α-linkages to fucose residues. The results showed that csFase I was highly active on α-1,2 linked fucolylated substrates (**Figure. 1**).

**Figure 1.**
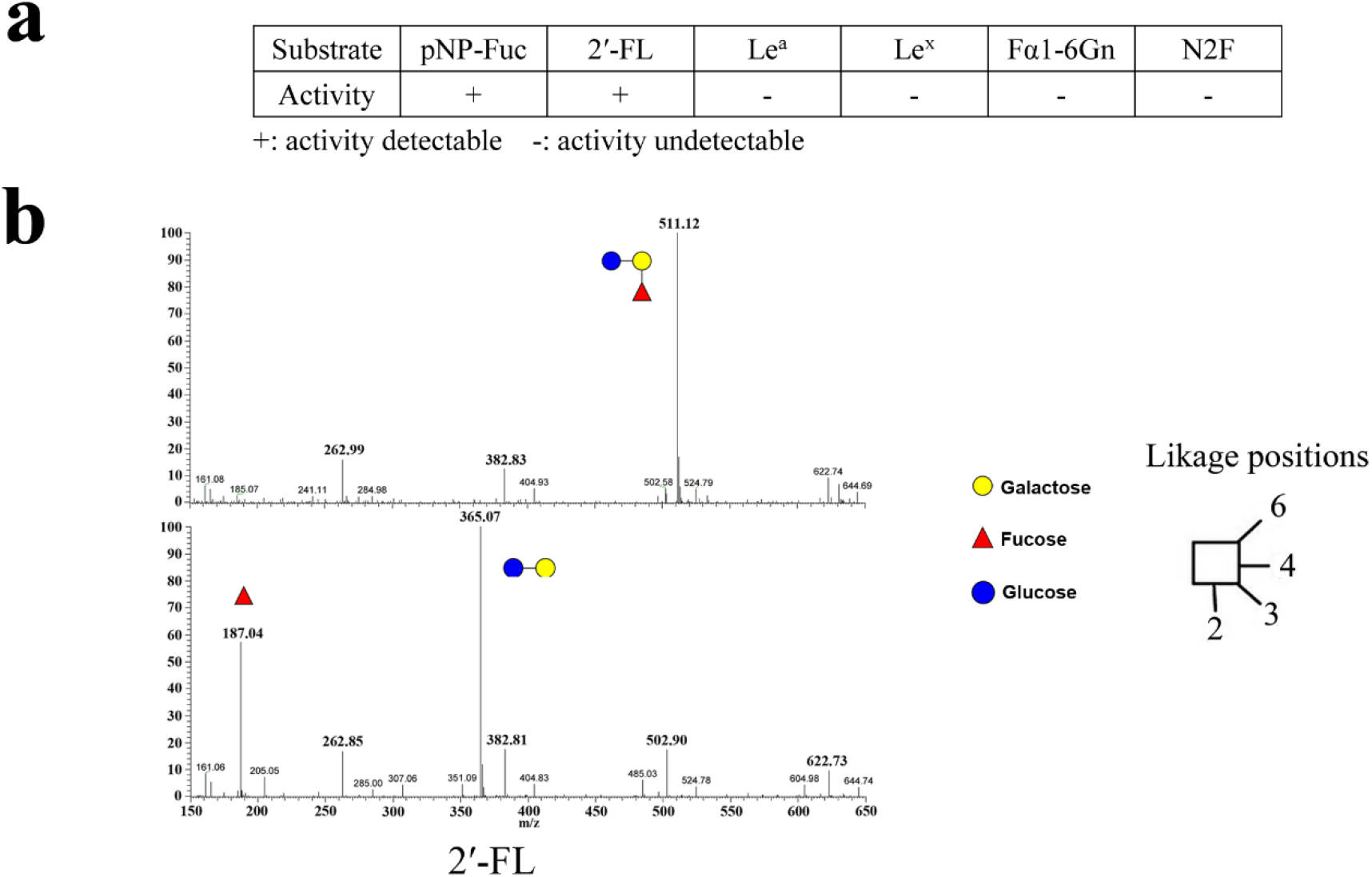
Determination of the linkage specificity of csFase I tested with a panel of fucosylated oligosaccharides. **a**) The enzymatic activity of csFase I against various fucosylated oligosaccharides was measured. **b**) The high activity of csFase I against 2’-FL were determined by MS analysis.

As ABO blood antigens contain α-1,2 linked fucose, we then measured the activity of the enzyme on blood group-associated oligosaccharides. The results showed that the enzyme is active only on the unbranched α-1,2 fucosylated H blood group, while no detectable activity against branched A and B oligosaccharides was detected under K-FUCOSE and MS detection (**Supplementary Figure 2, Supplementary Table S1**). The kinetic properties of the recombinant enzyme against H blood group substrates are presented in **Supplementary Table S2**, indicating its high enzymatic activity.

The fucosidase csFase I showed highly suitable kinetic properties for the removal of fucose from H blood group oligosaccharides. We then determined the activity of csFase I against RBCs with routine detection methods. Our results indicated that the complete removal of H antigens from O-type RBCs was achieved in isotonic phosphate buffer (pH 6.5) as evaluated by agglutination typing with clinically used reagents and methods (**Supplementary Figure 3**), as well as by sensitive laser confocal microscopy and fluorescence-activated cell sorting analysis (**Figure. 2a&b**, **Supplementary Figure 4**). In addition, treatment with csFase I could also completely remove H antigens from A-, B- and AB-type RBCs (**Fig. 1b**). Moreover, our results showed that csFase I was inactive against the fucose of A and B antigens on the surface of A, B or AB RBCs (**Supplementary Figure 3 b**), demonstrating its safety without the disruption of A and B antigens.

**Figure 2.**
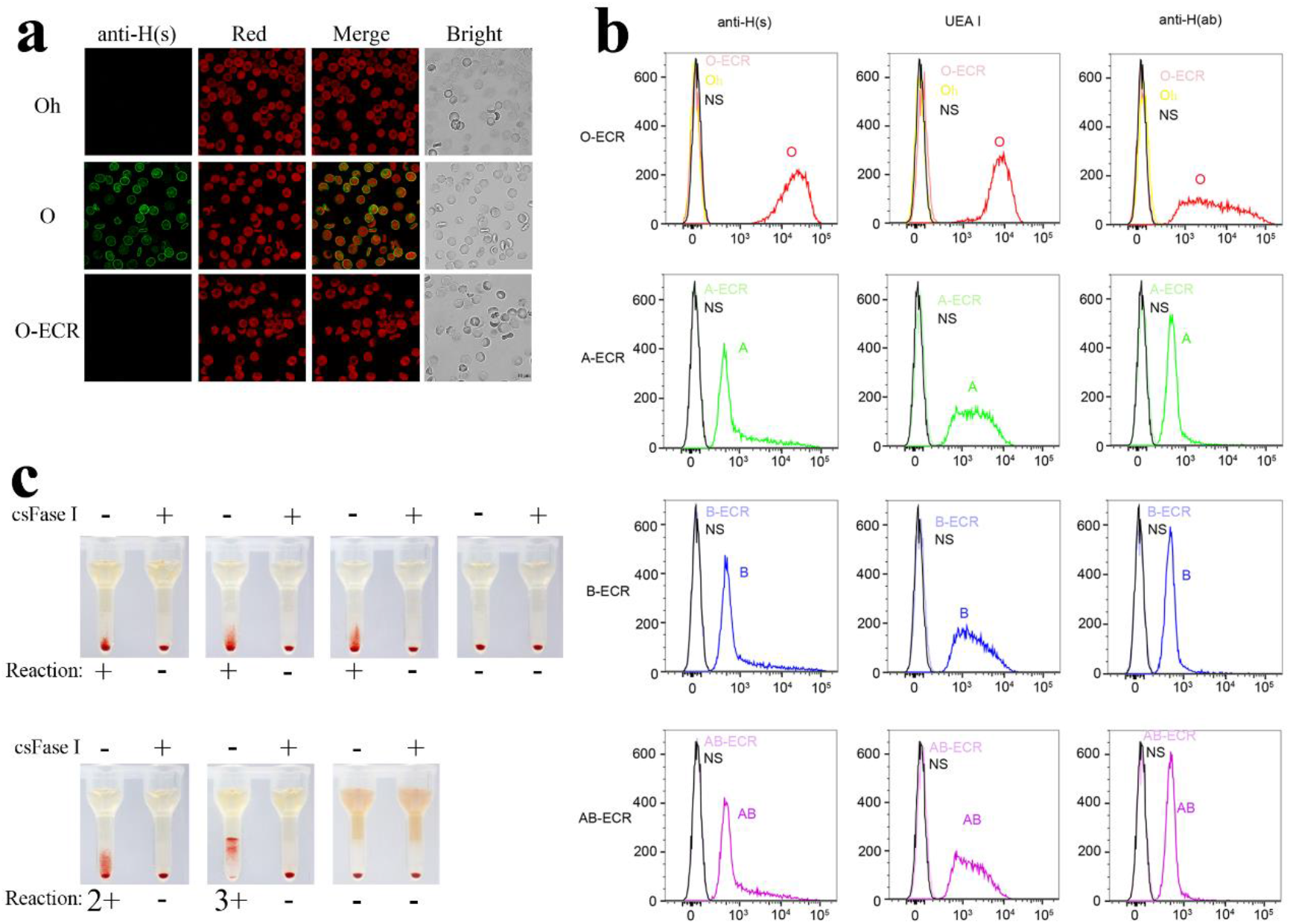
Complete removal of H antigens on the surface of RBCs by csFase I treatment. **a)** H antigens on the surface of native or csFase I-treated O RBCs were detected using immunofluorescence confocal microscopy. Oh: H-deficient RBC, negative control; O: O blood group RBC; O-ECR: O type of csFase I enzyme-converted RBC (ECR). **b)** FACS analysis of native and various ECRs. The x-axis represents the fluorescence intensity on a logarithmic scale, whereas the y-axis shows the number of RBCs evaluated. Oh: The yellow line of H-deficient RBCs in the O-ECR staining histogram (upper panels) represents the negative control, RBCs from a Para-Bombay donor lacking H antigens, as we did not find a Bombay-type donor. NS: The black line represents nonstaining RBCs, which also serve as a negative control. The red, green, blue and purple lines represent O, A, B and AB RBCs, respectively, and the line in the same slight color represents the corresponding ECR. The RBCs were stained with three anti-H grouping reagents from Shanghai SHPBC Corporation Limited (anti-H(s)), Sigma (UEA I) and Abcam (anti-H(ab)), respectively. **c)** Crossmatch of O-ECR with a panel of Para-Bombay sera. csFase I-treated or untreated O-type RBCs were subjected to a gel column and incubated for 15 min at room temperature. Then, the agglutination reaction was detected following the manufacturer’s protocol.

The buffer plays a major role in the enzymatic reaction of blood group conversion. To establish the optimum buffer conditions for fucosidase activity and simplify the conversion process, we measured the enzymatic activity of csFase I in routine RBC preservative solutions, such as MAP, SAGM, CPDA-1, Alsever’s solution and ACD-B^16,17^. The results showed that csFase I has higher activity in MAP, SAGM, CPDA-1 and Alsever’s solution than in citrate and phosphate buffers (**Supplementary Figure 5**).

We then measured the enzyme consumption of the conversion with a series of dose titrations in the optimum RBC preservative solutions. The results indicated that the completion of conversion has a low enzyme consumption of 84 μg/mL packed O type RBCs in 60 min of incubation (**Table 1**). Lower enzyme consumption was observed for the removal of H antigens from packed A, B and AB RBCs (**Supplementary Table S3**), as fewer H antigens are present on the surface of A-, B- and AB-type RBCs than on O-type RBCs.

**Table 1.** Blood group typing of O-ECR with enzyme dose titrations

| mg/mL<br>(pRBCs) | MAP |  |  | SAGM |  |  | CPDA-1 |  |  | Alsever's Solution |  |  |
| --- | --- | --- | --- | --- | --- | --- | --- | --- | --- | --- | --- | --- |
|  | anti-H(s) |  | UEA I | anti-H(s) |  | UEA I | anti-H(s) |  | UEA I | anti-H(s) |  | UEA I |
|  | 37°C | 4°C | RT | 37°C | 4°C | RT | 37°C | 4°C | RT | 37°C | 4°C | RT |
| 0.21 | 0 | 0 | 0 | 0 | 0 | 0 | 0 | 0 | 0 | 0 | 0 | 0 |
| 0.084 | 0 | 0 | 0 | 0 | 0 | 0 | 0 | 1+ | W+ | 0 | 0 | 0 |
| 0.042 | 0 | Vw+ | 0 | Vw+ | w+ | w+ | w+ | 2+ | 2+ | 0 | 1+ | 1+ |
| 0.021 | Vw+ | W+ | w+ | w+ | 1+ | 1+ | 1+ | 3+ | 2+ | 1+ | 2+ | 2+ |
| 0.0105 | W+ | 1+ | 1+ | 1+ | 2+ | 2+ | 2+ | 4+ | 3+ | 2+ | 3+ | 3+ |

Typing with anti-H reagents and methods as indicated using agglutination score 0 to 4+.
Enzyme-converted red blood cells were incubated at 37°C or 4°C for 15 min.
pRBCs, packed RBCs.
RT, room temperature.

Moreover, we collected a small panel of sera from Para-Bombay individuals to demonstrate the possibility of transfusion to a person with anti-H antibody. The results showed that five out of seven sera exhibited different levels of agglutination when incubated with O-type RBCs, while these agglutinations were completely eliminated when the O-type RBCs were treated with csFase I (**Figure. 2c**), demonstrating the great potential of csFase I in transfusion for the Bombay or Para-Bombay populations.

In summary, we report a potent and handy approach that enables the production of H antigen-deficient RBCs for transfusion. Our approach may allow improvement of the blood supply and enhancement of patient safety in transfusion medicine, especially for the rare population with the Bombay and Para-Bombay phenotypes.

## Materials and Methods

### Materials and Reagents

All p-nitrophenyl (pNP-) monosaccharides and fucosylated oligosaccharides used in this study were obtained from Carbosynth (Compton, U. K), except for the blood-associated oligosaccharides, which were purchased from Elicityl OligoTech (Crolles, France). Free fucose was assayed with the K-FUCOSE kit from Megazyme International (Dublin, Ireland). Murine monoclonal anti-A and anti-B blood grouping reagents were obtained from Changchun Brother Biotech Corporation Limited Changchun, China. Anti-H antibodies were obtained from Shanghai SHPBC Corporation Limited and Abcam, respectively. Ulex Europaeus Agglutinin I (UEA I) and FITC-conjugated UEA I were purchased from Sigma. Alex fiur-488 conjugated anti-mouse IgM was obtained from Abcam. Red blood cells from normal blood donors with the ABO phenotype were obtained from Shanghai Blood Centre. The sera from Para-Bombay type donors were collected by Fujian Union Hospital, and the usage of the sera for this study was approved by the Ethics Committee of the hospital.

### Enzyme assays

Assays with chromogenic pNP substrates were performed in the isotonic phosphate buffer at pH 6.5, except for measurement of the optimal pH, as previously described. Reactions were terminated by the addition of 1.5-fold reaction volumes of 1 M Na_2_CO_3_, and pNP formation was quantified at 405 nm. Assays with oligosaccharide structures were carried out in a reaction mixture of 10 μL. Product formation was measured by the K-Fucose Kit and electrospray ionization-mass spectrometry (ESI-MS) following the manufacturer’s instructions.

To determine the values of kinetic constants, 0.05 to 4 mM pNP-α-L-Fuc (pNP-Fuc) or oligosaccharide was coincubated with a constant amount of each enzyme. The time course for each reaction was previously determined to fulfill steady-state assumptions, and product formation was determined as mentioned above. Nonlinear regression was used to determine K_m_ and V_max_ values by using the program GraphPad Prism software version 5.0 (GraphPad Software, San Diego, California, USA). All reactions were performed in triplicate for statistical evaluations.

### ESI-MS

Electrospray ionization-mass spectrometry, which was used to determine the hydrolysis activity against each substrate, was conducted on a TSQ Quantiva™ triple quadrupole mass spectrometer system (Thermo Fisher Scientific, MA, USA). A 10 μL reaction mixture sample was diluted in 200 μL of 18.2 MΩ ultrapure water and injected into a mass spectrometer at 20 μL/min with a microsyringe pump (Pump 22; Harvard Apparatus, MA). The instrument was operated in a full scan model, from m/z 100 to 900 for 2 min in positive mode.

### Enzymatic conversion of RBCs with fucosidase

Enzymatic conversion reactions were performed mainly based on a previous study. Briefly, fresh whole blood was obtained from Shanghai Blood Center, and the buffy coat was removed. The packed red blood cells were prepared by prewashing in conversion buffer 3 times. The enzymatic conversion reaction was incubated for 60 min with gentle mixing at 26°C with a final 30% hematocrit of red blood cells. Then, the RBCs were washed four times with 1:4 vol/vol of saline by centrifugation at 500×*g* to remove the enzyme. The washed enzyme-converted RBCs were typed according to standard blood banking techniques by the routine method and sensitive FACS.

### Laser confocal microscopy

The erythrocytes were fixed and stained as previously described with minor changes^18^. Briefly, the cells were washed twice in PBS and then fixed for 10 min in PBS containing 0.5% glutaraldehyde. The cells were then rinsed four times with PBS to remove the unreacted glutaraldehyde. Fixed RBCs were stained by using antibodies or reagent with appropriate dilution or undiluted for 20 min. Specifically, the monoclonal anti-H(ab) antibody from Abcam (1:1000), FITC labeled UEA I (2 μg/mL) from Sigma, and anti-H(s) antibody from SHPBC (undiluted) were used to stain the indicated RBCs for 20 min. After washing three times with PBS to remove the unreacted antibodies, the cells were again labeled with secondary antibody. After labeling, resuspended RBCs were allowed to attach to cover slips coated with polylysine, and the cover slips were mounted using M800 Aqua Mount and sealed with nail polish. The prepared samples were scaled with a Leica TCS SP8 confocal platform.

### Flow cytometry

The native and enzyme-treated RBCs were analyzed by flow cytometry. Briefly, 50 μL of 0.5% RBCs was pelleted and suspended in 50 μL PBS and then fixed with 0.1% glutaraldehyde in PBS for 10 min, followed by three washes in PBS. Cells were stained with first and second antibodies as mentioned above in confocal microscopy detection. Finally, the stained cells were resuspended in 500 μL of PBS for further flow cytometry analysis with a FACSCalibur system (Becton, Dickinson and Company, Franklin Lakes, NJ, USA).

## ACKNOWLEDGMENTS

We gratefully acknowledge Dong Xiang for assistance with blood typing and discussion. We also thank Zebin Lin for help with mass spectrometry. This work was funded in part by the National Science and Technology Major Project (2014ZX09101046-004) to L.C and the National Natural Science Foundation of China (31600644, 31801157) to G.Q.S and J.Y.

## COMPETING INTERESTS

The authors declare no competing interests.

