## Supplemental Figures&Tables for "Bacterial fucosidase enables the production of Bombay red blood cells"

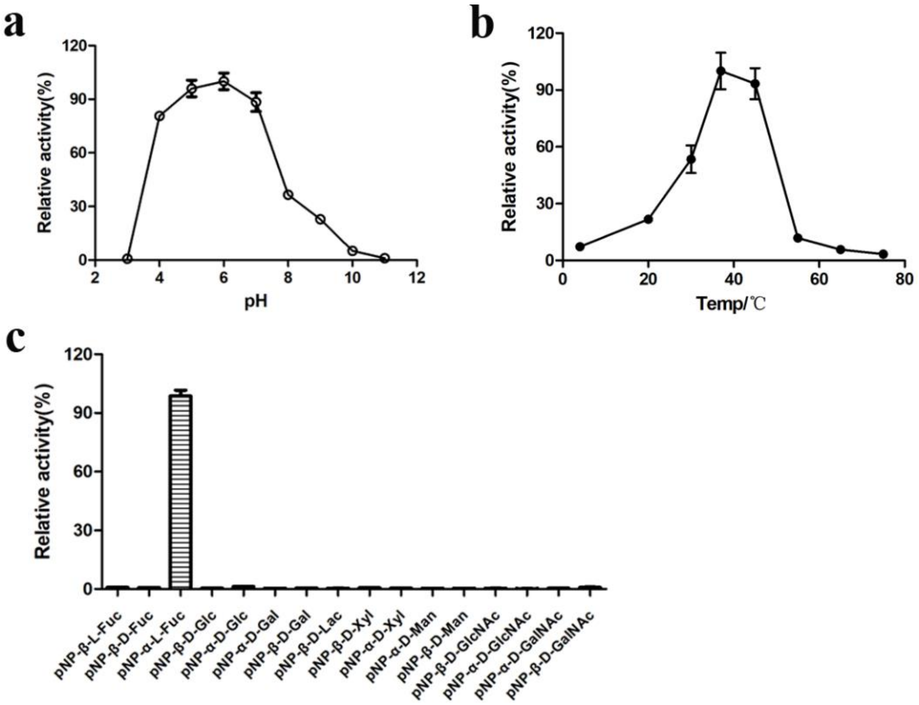


Supplementary Figure 1

The characteristics of csFase I. **a&b**) The enzymatic activity of csFase I at various pH values and temperatures shows an optimum pH between 5 and 7 and an optimum temperature of 37°C for csFase I. **c**) Substrate specificity of cFase I against various pNP-monosaccharides. The substrate was used at a concentration of 1.0 mM. The activity on pNP-α-L-Fuc was taken to be 100%.

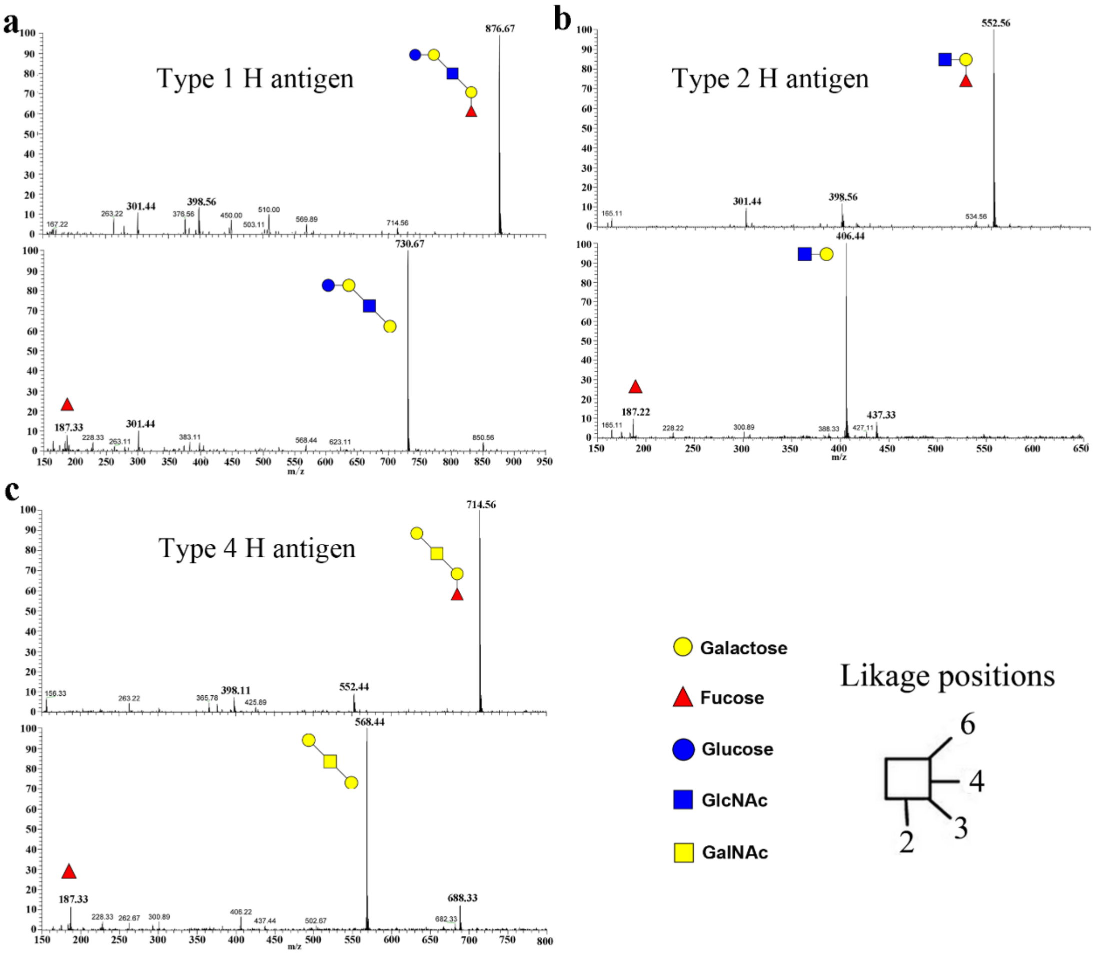


Supplementary Figure 2

The activity of csFase I against blood group-associated oligosaccharides. type 1 (**a**), type 2 (**b**) and type 4 (**c**) H antigens of blood group-associated oligosaccharides were subjected to csFase I degiestion, respectively, and product formations were determined by MS analysis. The results indicated that csFase I was highly active against the blood group-associated oligosaccharides.

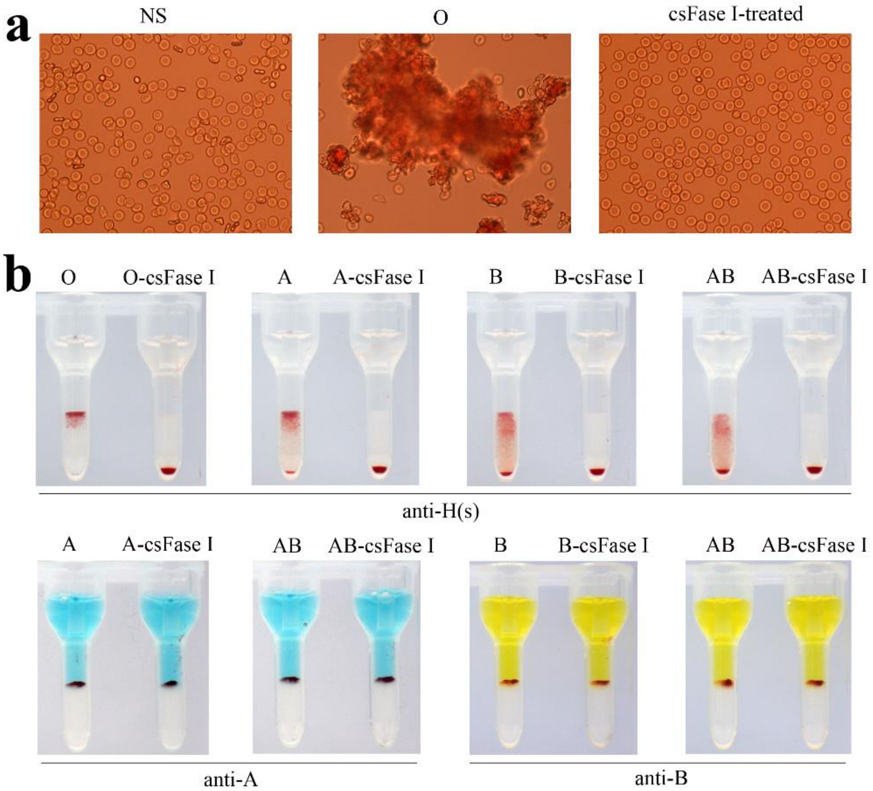


Supplementary Figure 3

The removal of H antigens from RBCs was detected by routine methods. **a**) The enzymatic activity of csFase I against H antigens of O-type RBCs was determined by the conventional slide method. NS, nonstaining; **b**) The enzymatic activity of csFase I against the H antigens of O-, A-, B- and AB-type RBCs was determined (upper panel), and no activity of csFase I against the branched fucose of A and B antigens was observed (lower panel) by card agglutination.

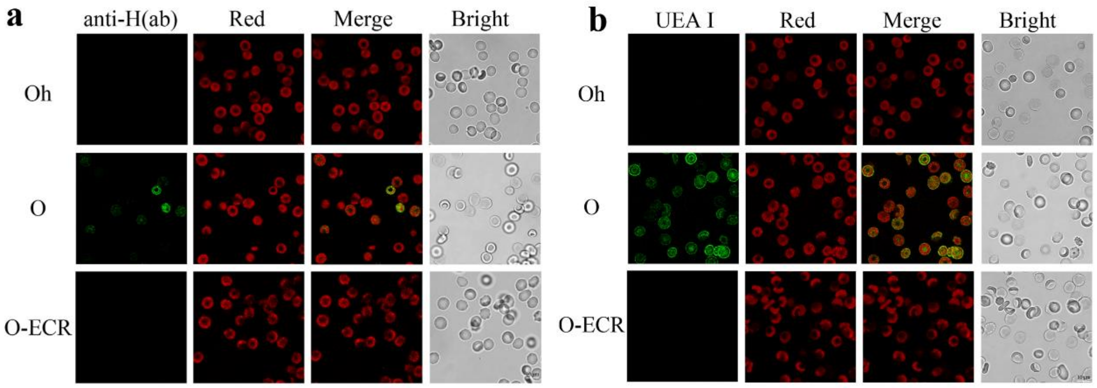


Supplementary Figure 4

Complete removal of H antigens on the surface of O-RBCs by csFase I treatment. H antigens on the surface of native or csFase I-treated O-RBCs were detected with anti-H(ab) antibody (**a**) and FITC-UEA I (**b**) by immunofluorescence confocal microscopy.

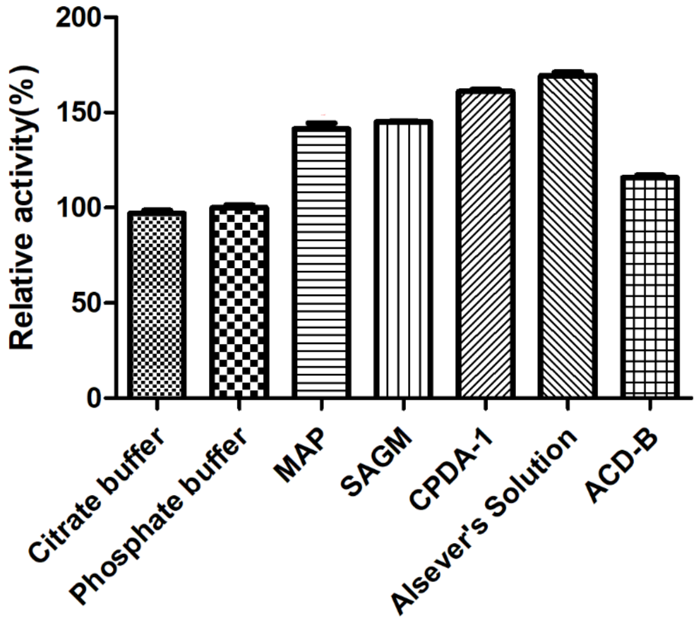


Supplementary Figure 5

The enzymatic activity of csFase I in routine RBC preservative solutions. A higher activity in MAP, SAGM, CPDA-1 and Alsever's solution was observed compared with that in the citrate and phosphate buffers.

Supplementary Table S1. Substrate specificities analyzed by K-FUCOSE kit and MS

| Blood group specificity | Substrate | Activity | |
|---|---|---|---|
| | | K-FUCOSE | MS |
| **Blood group A structure** | | | |
| type 1 tetraose | GalNAcα-3(Fucα-2)Galβ-3GlcNAc | - | - |
| type 2 hexaose | GalNAcα1-3(Fucα1-2)Galβ1-4GlcNAcβ1-3-Lac | - | - |
| type 5 tetraose | GalNAcα1-3(Fucα1-2)Galβ1-4Glc | - | - |
| **Blood group B structure** | | | |
| type 1 tetraose | Galα1-3(Fucα1-2)Galβ1-3GlcNAc | - | - |
| type 2 hexaose | Galα1-3(Fucα1-2)Galβ1-4GlcNAcβ1-3-Lac | - | - |
| type 5 tetraose | Galα1-3(Fucα1-2)Galβ1-4Glc | - | - |
| **Blood group H structure** | | | |
| type 1 pentaose | Fucα1-2Galβ1-3GlcNAcβ1-3Galβ1-4Glc | + | + |
| type 2 triaose | Fucα1-2Galβ1-4GlcNAc | + | + |
| type 4 tetraose | Fucα1-2Galβ1-3GalNAcβ1-3Gal | + | + |
| type 5 triaose (2'-FL) | Fucα1-2Galβ1-4Glc | + | + |
| H disaccharide | Fuca1-2Gal | + | + |

-: activity undetectable +: activity detectable

Supplementary Table S2. Kinetic properties of csFase I on pNP-Fuc and blood group H oligosaccharides.

| Substrate | $k_{cat}(s^{-1})$ | $K_m$(mM) | $k_{cat}/K_m(s^{-1}/mM)$ |
|---|---|---|---|
| pNP-Fuc | 0.13±0.002 | 0.33±0.02 | 0.39±0.03 |
| H disaccharide | 13.05±1.25 | 1.22±0.08 | 10.69±1.26 |
| type 1 | 26.69±2.52 | 0.72±0.09 | 36.82±5.91 |
| type 2 | 30.50±1.71 | 2.66±0.28 | 11.45±1.37 |
| type 4 | 121.82±5.66 | 2.74±0.24 | 44.49±4.39 |
| type 5 | 32.34±2.43 | 0.70±0.05 | 46.09±4.83 |

Supplementary Table S3. Blood group typing of A, B and AB-ECR with enzyme dose titrations

| | | MAP | | | SAGM | | | CPDA-1 | | | Alsever's Solution | | |
|---|---|---|---|---|---|---|---|---|---|---|---|---|---|
| | mg/mL | anti-H(s) | | UEA I | anti-H(s) | | UEA I | anti-H(s) | | UEA I | anti-H(s) | | UEA I |
| ECRs | (pRBCs) | 37℃ | 4℃ | RT | 37℃ | 4℃ | RT | 37℃ | 4℃ | RT | 37℃ | 4℃ | RT |
| A-RBC | | | | | | | | | | | | | |
| | 0.084 | 0 | 0 | 0 | 0 | 0 | 0 | 0 | 0 | 0 | 0 | 0 | 0 |
| | 0.042 | 0 | 0 | 0 | 0 | 0 | 0 | 0 | 0 | 0 | 0 | 0 | 0 |
| | 0.021 | 0 | 0 | 0 | 0 | 0 | 0 | W+ | 1+ | Vw+ | 0 | 0 | 0 |
| | 0.0084 | 0 | W+ | 0 | Vw+ | W+ | 0 | 1+ | 2+ | W+ | 0 | 1+ | W+ |
| | 0.0042 | W+ | 1+ | W+ | W+ | 1+ | W+ | 1+ | 3+ | 1+ | W+ | 2+ | 1+ |
| | 0.0021 | 1+ | 2+ | 2+ | 1+ | 2+ | 1+ | 2+ | 4+ | 2+ | 2+ | 3+ | 2+ |
| B-RBC | | | | | | | | | | | | | |
| | 0.084 | 0 | 0 | 0 | 0 | 0 | 0 | 0 | 0 | 0 | 0 | 0 | |
| | 0.042 | 0 | 0 | 0 | 0 | 0 | 0 | 0 | W+ | Vw+ | 0 | 0 | 0 |
| | 0.021 | 0 | 0 | 0 | 0 | Vw+ | 0 | W+ | 1+ | W+ | 0 | 0 | 0 |
| | 0.0084 | Vw+ | W+ | 0 | Vw+ | W+ | W+ | 1+ | 2+ | 1+ | 0 | 1+ | 1+ |
| | 0.0042 | W+ | 1+ | W+ | W+ | 1+ | 1+ | 1+ | 3+ | 2+ | W+ | 2+ | 2+ |
| | 0.0021 | 1+ | 2+ | 1+ | 1+ | 2+ | 2+ | 2+ | 3+ | 3+ | 2+ | 3+ | 3+ |
| AB-RBC | | | | | | | | | | | | | |
| | 0.084 | 0 | 0 | 0 | 0 | 0 | 0 | 0 | 0 | 0 | 0 | 0 | 0 |
| | 0.042 | 0 | 0 | 0 | 0 | 0 | 0 | 0 | Vw+ | 0 | 0 | 0 | 0 |
| | 0.021 | 0 | 0 | 0 | 0 | 0 | 0 | 0 | W+ | W+ | 0 | Vw+ | 0 |
| | 0.0084 | 0 | W+ | 0 | 0 | 0 | 0 | 0 | 1+ | 1+ | 0 | W+ | W+ |
| | 0.0042 | Vw+ | 1+ | W+ | 0 | 1+ | 1+ | 1+ | 2+ | 2+ | 0 | 1+ | 1+ |
| | 0.0021 | W+ | 1+ | 1+ | W+ | 2+ | 2+ | 2+ | 3+ | 3+ | W+ | 2+ | 2+ |

Typing with anti-H reagents and methods as indicated using agglutination score 0 to 4+.
Enzyme-converted red blood cells were incubated at 37°C or 4°C for 15 min.
pRBCs, packed RBCs.
RT, room temperature.
